## Supplementary material for "Heterochromatin *de novo* formation and maintenance in *Plasmodium falciparum*": Sup. Fig. S1-10 and Sup. Table 1

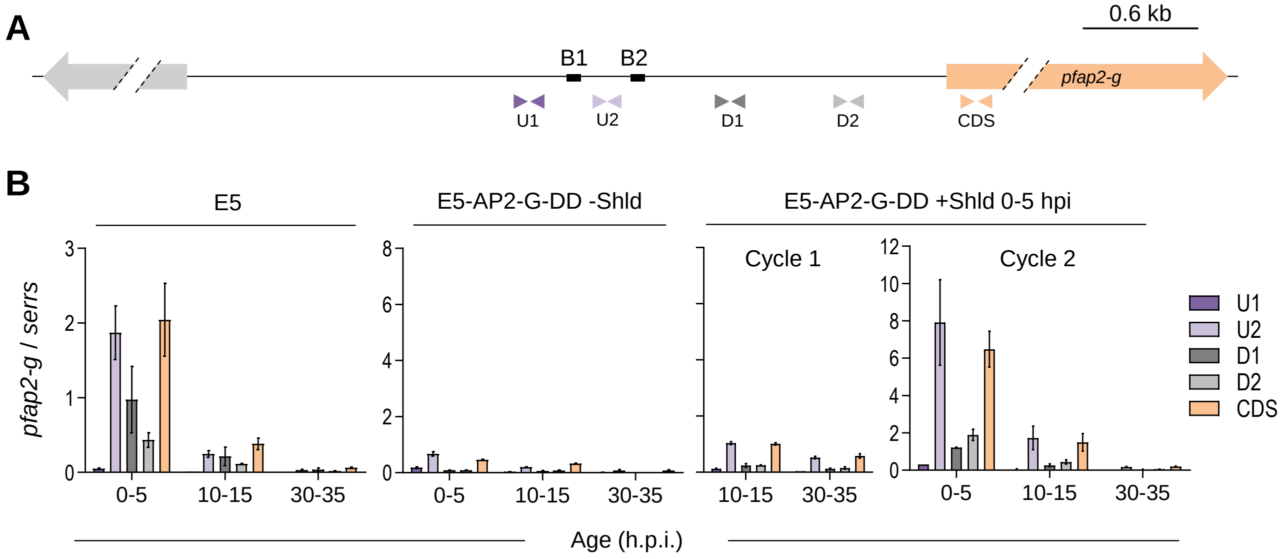


**Fig. S1. RT-qPCR validation of the *pfap2-g* TSSs**

**(A)** Schematic of the position of the primers (colored arrowheads) used for validation of the TSSs identified. The position of TSSs block 1 (B1) and block 2 (B2) is also shown.

**(B)** Validation of the TSSs by RT-qPCR with samples from tightly synchronized E5 and E5-AP2-G-DD cultures, the latter prepared either in the absence (-Shld) or presence (+Shld) of Shld1 and analyzed at the cycle of stabilization (Cycle 1) or at the next cycle (Cycle 2). In cultures with Shld1, the compound was added at 0-5 h post-invasion (hpi) of Cycle 1. Transcript levels are normalized against serine-tRNA ligase (*serrs*). Values are the average of two biological replicates, with s.e.m.

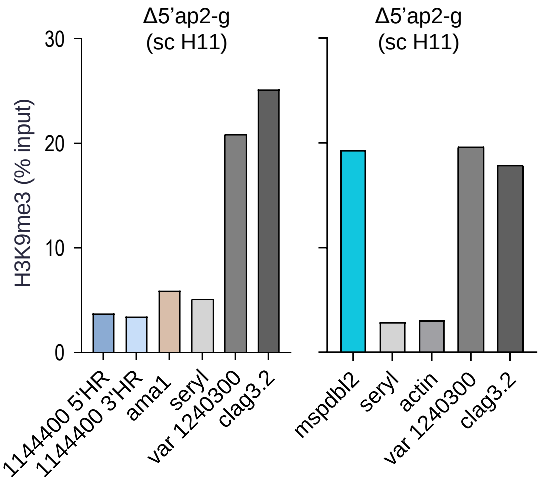


**Fig. S2. Validation of the euchromatic or heterochromatic state of genes analyzed in this study in the Δ5'ap2-g line**

H3K9me3 ChIP-qPCR coverage, calculated as the % DNA recovered in the H3K9me3 IP relative to the input, in the Δ5'ap2-g line (subclone H11). The genes analyzed were the PF3D7_1144400 gene (1144400 5'HR and 3'HR) used for fragment integration, euchromatic genes *ama1* (PF3D7_1133400), serine-tRNA ligase (*serrs*; PF3D7_0717700) and actin I (*actI*; PF3D7_1246200), and heterochromatic genes *var* 1240300 (PF3D7_1240300), *clag3.2* (PF3D7_0302200) and *mspdbl2* (PF3D7_1036300). In the HC nucleation and maintenance ChIP-qPCR experiments, the genes *serrs* and *actI* were used as ChIP-qPCR negative controls, whereas the genes *var* 1240300 (PF3D7_1240300), *clag3.2* and *mspdbl2* were used as ChIP-qPCR positive controls.

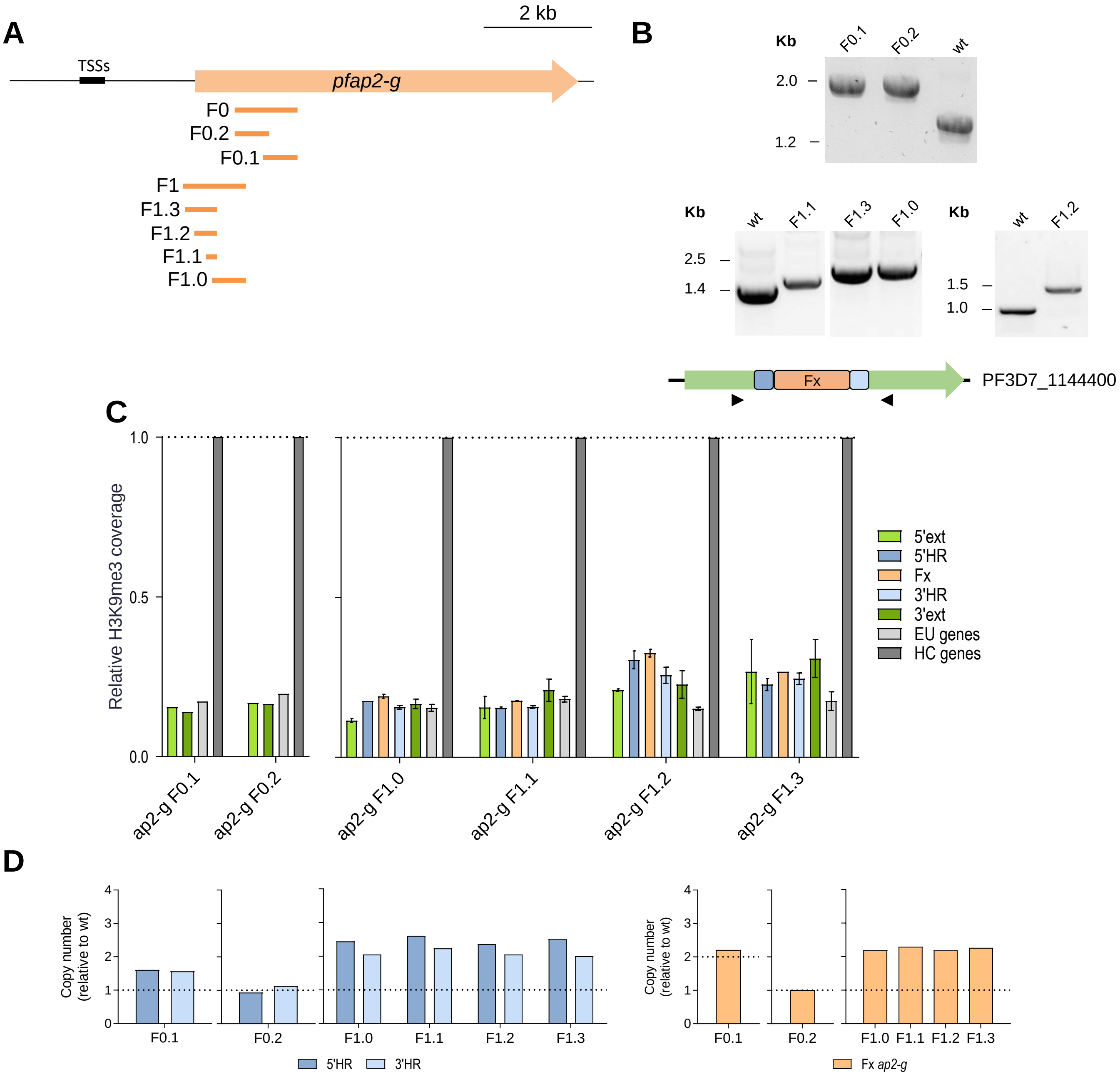


**Fig. S3. Assessment of HC nucleation by small *pfap2-g* fragments derived from fragments F0 and F1**

**(A)** Schematic (to scale) of the position of the small *pfap2-g* fragments derived from the F0 and F1 fragments.

**(B)** Diagnostic PCR confirming the correct integration of each fragment in the transgenic lines. The position of the PCR primers, external to the HRs, are shown in the scheme at the bottom.

**(C)** H3K9me3 ChIP-qPCR analysis of the transgenic lines carrying small (<1kb) *pfap2-g* fragments derived from *pfap2-g* F0 and F1 fragments. The position of the primers is as in Fig. 2A. Values are the % of the DNA recovered in the H3K9me3 IP (% input) at each position relative to the average % input in the positive control heterochromatic genes *var* PF3D7_1240300 and *clag3.2* (“HC genes”). “EU genes” are euchromatic genes *actI* and *serrs* used as negative controls (average of the two shown). The dashed line indicates the level of enrichment in the positive control heterochromatic genes. Data are presented as the average and s.e.m. of two biological replicates (except for F0.1 and F0.2, N=1).

**(D)** qPCR analysis of the number of copies of the PF3D7_1144400 HRs (5’HR and 3’HR) and the fragments under analysis (Fx) in the gDNA of the transgenic lines. Copy number was calculated relative to 1.2B (wild type) gDNA. The dashed lines indicate the expected value for single, correct integration.

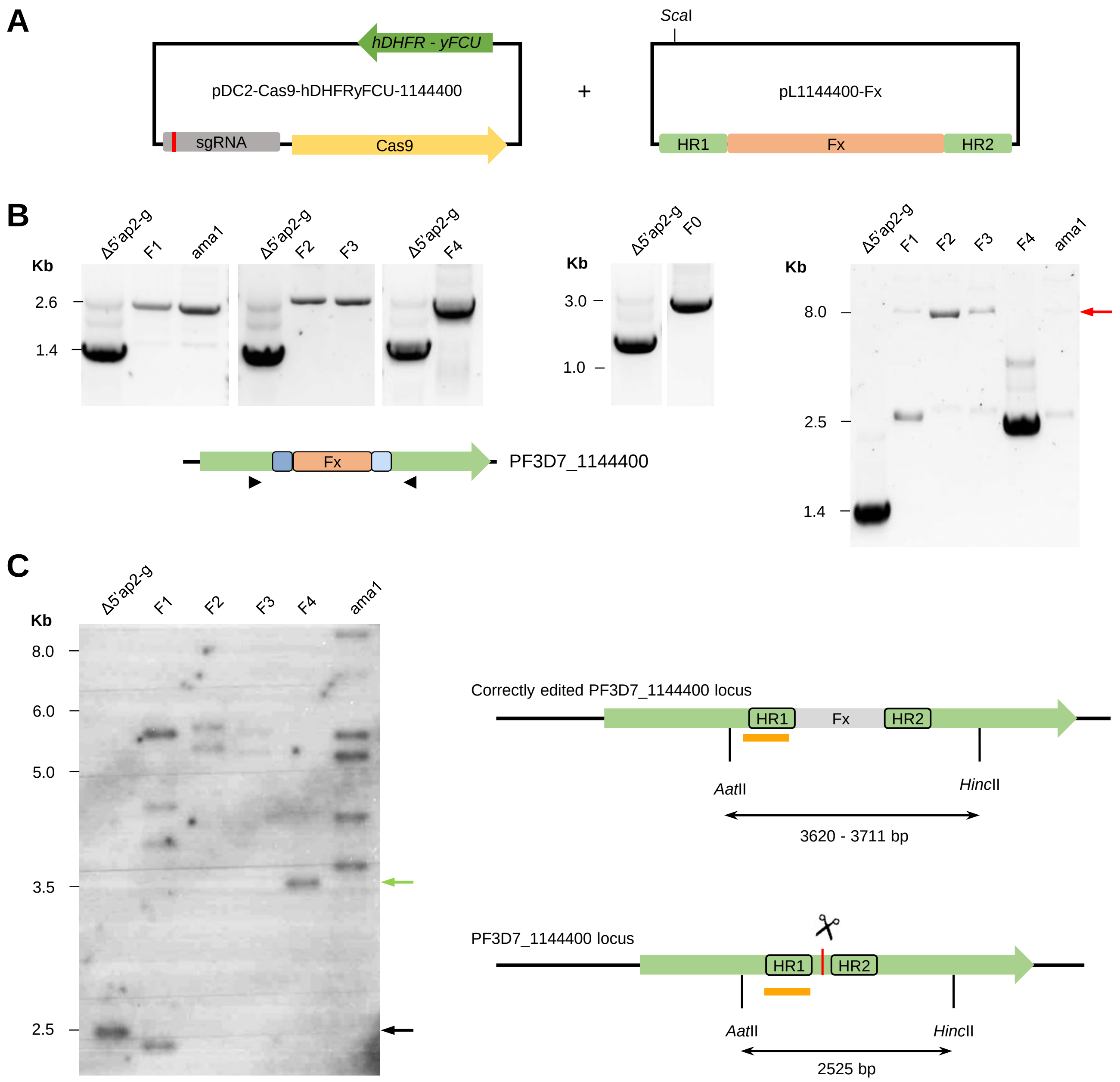


**Fig. S4. Integration of ~1 kb fragments into the PF3D7_1144400 locus**
**(A)** Schematic (not to scale) of the plasmids used to generate the transgenic lines carrying different *pfap2-g* ~1 kb fragments (F0 to F4) and an *ama1* fragment. Note that the *var* 1240300 transgenic line was generated with the plasmids described in Fig.S9 (containing the *yfcu* negative selection marker) but for the experiments presented in Fig. 2 it was analyzed before negative selection, which makes it analogous to the other transgenic lines in those experiments.
**(B)** Diagnostic PCR confirming the correct integration of each fragment in the transgenic lines. The position of the PCR primers, external to the HRs, is shown in the scheme at the bottom. The panel at the right is a PCR amplification with the same primers using a longer extension time (10 min), which in most samples revealed a band that likely corresponds to integration of concatemers (red arrow).

**(C)** Southern blot analysis of the parental Δ5'ap2-g line (H11 subclone) and the transgenic lines with integrated fragments *pfap2-g* F1-F4 and ama1. The schematic shows the expected size of gDNA digested with *Aat*II, *Hinc*II and *Afl*II (the latter cleaving the plasmid but not the PF3D7_1144400 locus) and hybridized with a PF3D7_1144400-specific probe (orange line) in parasites with the fragments (Fx) correctly integrated (single copy) and in wild type parasites. The scissors indicate the position targeted by the guide RNA, whereas black boxes indicate the position of the HRs. The green arrow indicates the approximate position of the bands expected for the correctly edited locus, and the black arrow the position of the band expected for the wild type locus. While the F4 line showed the expected band for correct single integration, other lines showed a complex pattern consistent with integration of different plasmid concatemer species.

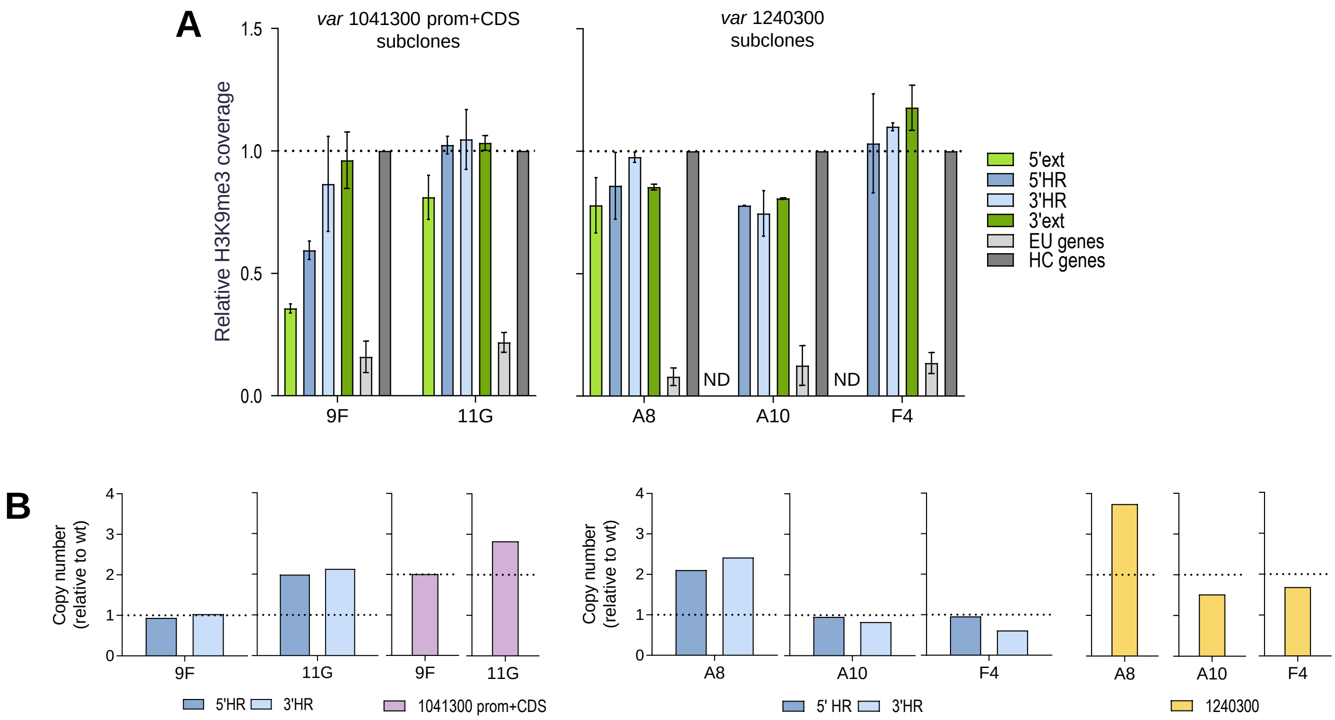


**Fig. S5. Assessment of HC nucleation in subclones of parasite lines carrying *var* fragments**

**(A)** H3K9me3 ChIP-qPCR analysis, as in Fig. S3C, of subclones derived from two transgenic lines carrying *var* fragments (1041300 prom+CDS and 1240300). ND indicates "not detected", as some subclones carried a spontaneous deletion at that position. However, ectopic HC formation was observed in the subclone not carrying the deletion as well as in subclones carrying the deletion. Data are presented as the average and s.e.m. of two biological replicates.

**(B)** qPCR analysis of copy number, as in Fig. S3D. The 1041300 prom+CDS subclones 9F and 11G carry single or multiple fragment integrations, respectively, but ectopic HC nucleation was observed in both subclones at similar levels.

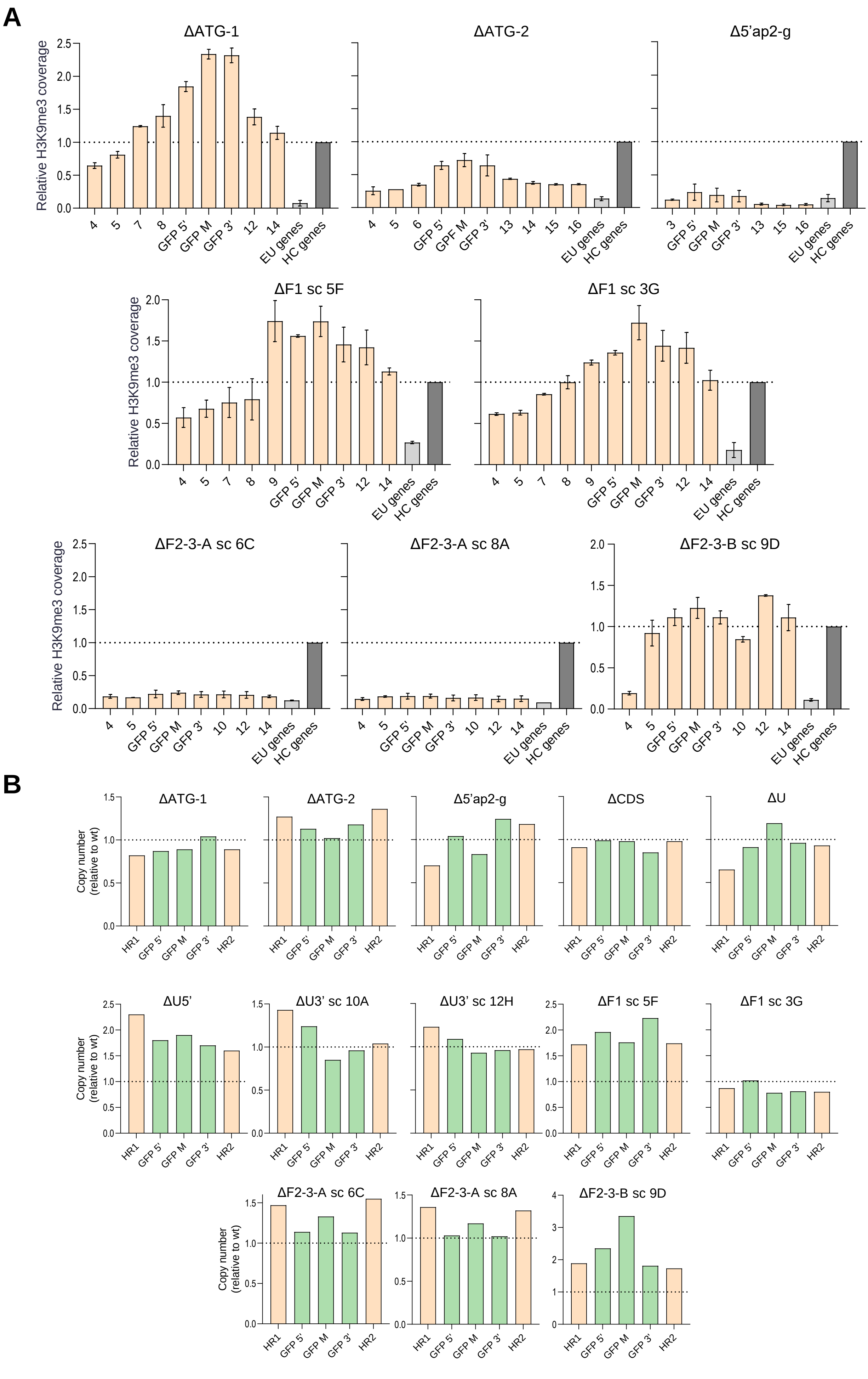
A
ΔATG-1
ΔATG-2
Δ5'ap2-g
Relative H3K9me3 coverage
0.0
0.5
1.0
1.5
2.0
2.5
4
5
7
8
GFP 5'
GFP M
GFP 3'
12
14
EU genes
HC genes
6
GPF M
13
15
16
3
ΔF1 sc 5F
ΔF1 sc 3G
9
ΔF2-3-A sc 6C
ΔF2-3-A sc 8A
ΔF2-3-B sc 9D
10
B
ΔCDS
ΔU
Copy number (relative to wt)
HR1
HR2
ΔU5'
ΔU3' sc 10A
ΔU3' sc 12H
1
2
3
4

**Fig. S6. Analysis of the role of different regions of *pfap2-g* in HC maintenance**

**(A)** H3K9me3 ChIP-qPCR analysis of the ΔATG-1, ΔATG-2 and Δ5'ap2-g transgenic lines, and subclones (sc) from the ΔF1 and ΔF2-3 lines. Values are the % input at each position relative to the % input in the positive control heterochromatic genes, as in Fig. S3C. Numbers in the x axis are the primer pairs shown in Fig. 7A. The dashed line indicates the level of enrichment in the positive control heterochromatic genes. Data are presented as the average and s.e.m. of two biological replicates.

**(B)** qPCR analysis of copy number, as in Fig. S3D, of the transgenic lines in panel A and in Fig. 6. Copy number was determined for the different *pfap2-g* HRs used for each line and for three different regions of the inserted gfp*. The dashed lines indicate the expected value for correct, single integration.

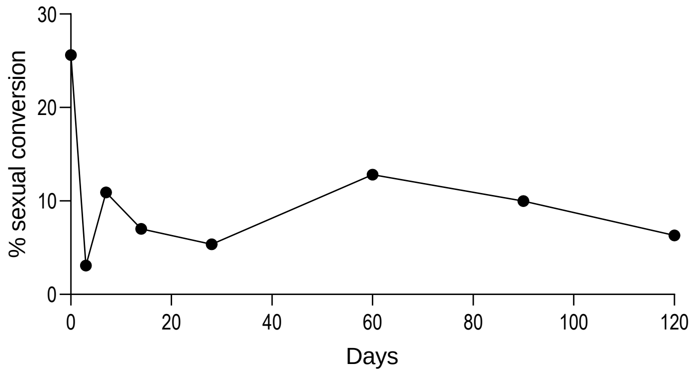


**Fig. S7. Evolution of the sexual conversion rate of E5-AP2-G-DD after a second exposure to Shld1 at different times**

Sexual conversion rate of E5-AP2-G-DD after induction with Shld1 for the first time (first data point, day 0), or after a second induction with Shld1 at different time points (cultures were maintained without Shld1 between the first and second exposures).

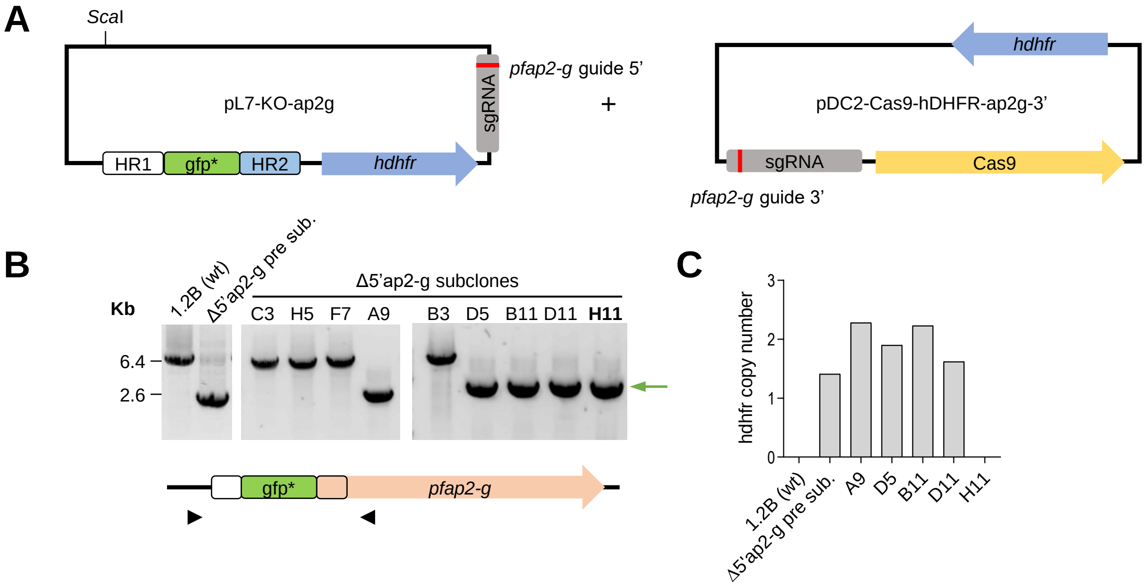


### Fig. S8. Generation of the Δ5'ap2-g line

**(A)** Schematic (not to scale) of the plasmids used to generate the Δ5'ap2-g line.
**(B)** Diagnostic PCR analysis of the parental 1.2B line (1.2B wt), the Δ5'ap2-g line before subcloning and its subclones. The position of the band expected for correctly edited parasites is indicated by a green arrow. The schematic shows the position of the PCR primers, external to the HRs.
**(C)** qPCR analysis of copy number of the *hdhfr* selectable marker. Quantification was performed against a standard curve prepared with gDNA of the W4-2 line (see Methods) previously shown by Southern blot to contain a single copy of the *hdfhr* gene integrated. Only the H11 clone carried the correct integration and was free of the marker and therefore was selected for all further experiments.

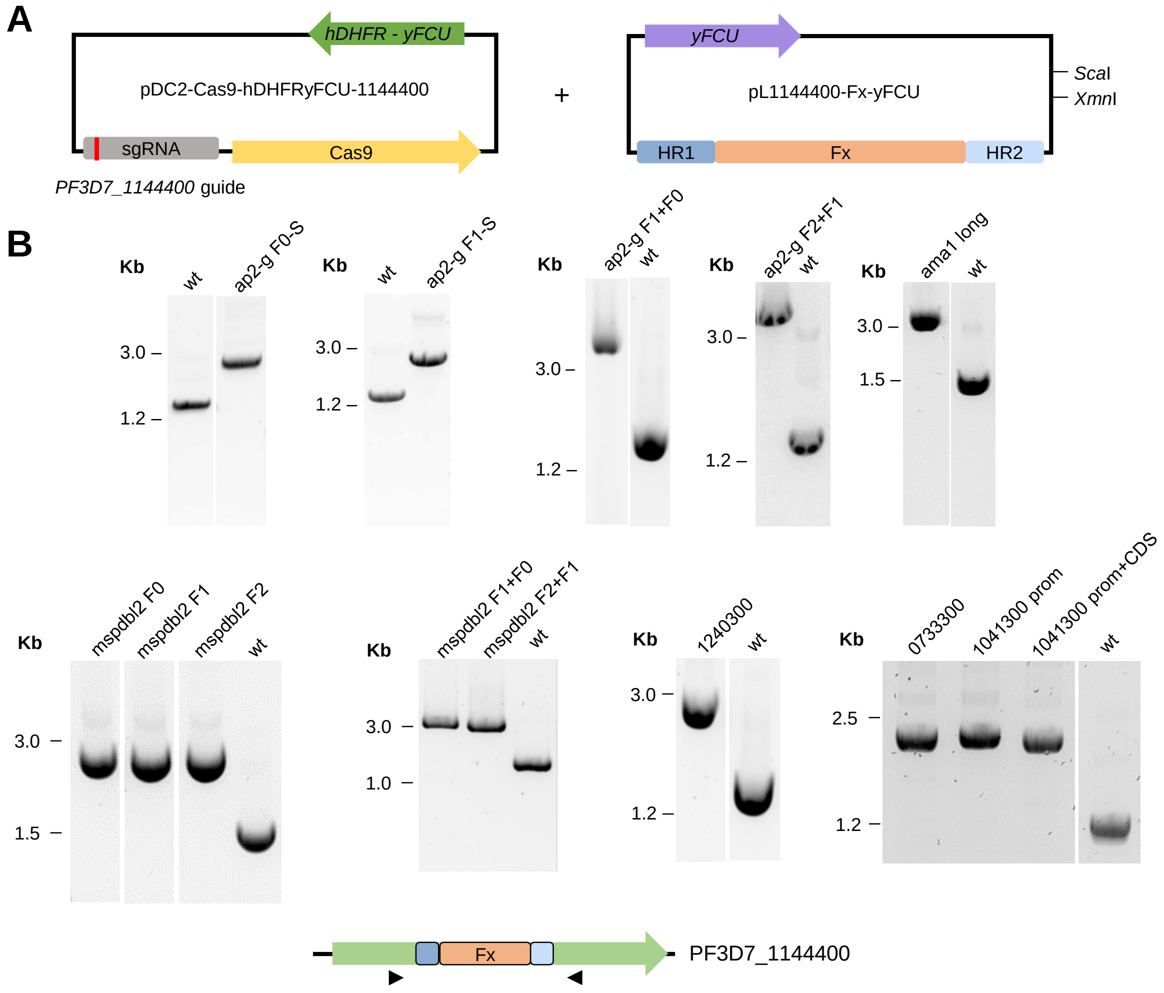


**Fig. S9. Generation of transgenic lines to assess HC nucleation by single copies of integrated fragments**

**(A)** Schematic (not to scale) of the plasmids used for the generation of transgenic lines with single copy fragment integrations, containing the *yFCU* negative selection marker.

**(B)** Diagnostic PCR of the transgenic lines generated using this strategy. The scheme at the bottom shows the position of the PCR primers, external to the HRs.

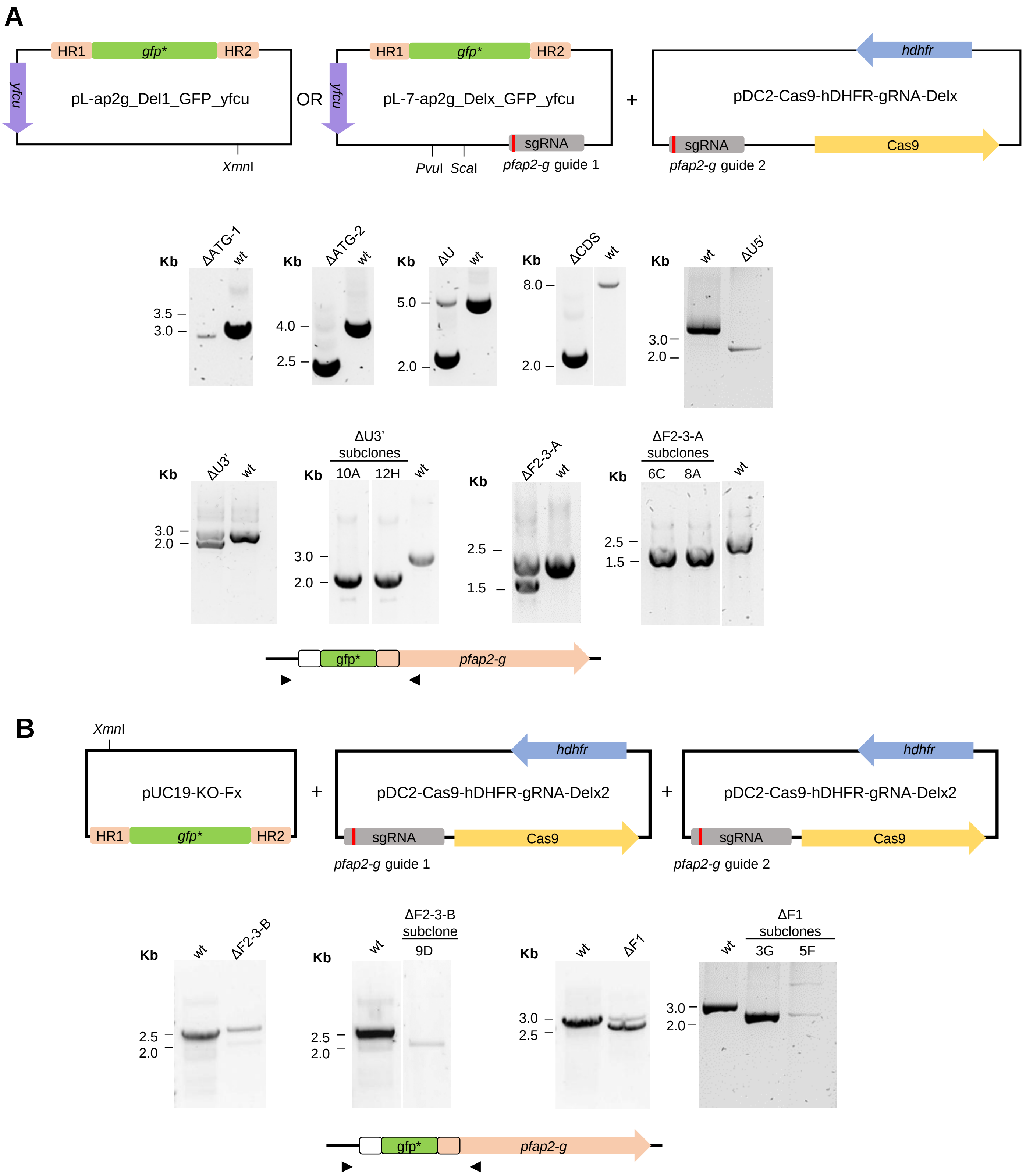


**Fig. S10. Generation of the transgenic lines to study HC maintenance at the *pfap2-g* locus**

**(A)** Top: Schematic (not to scale) of the plasmids used to delete different regions of the *pfap2-g* locus with a two plasmids strategy. The donor plasmid either lacked (ΔATG-1 line) or included (all other lines) a sgRNA expression cassette.

Bottom: Diagnostic PCR to assess correct edition in the transgenic lines generated with this strategy. The scheme at the bottom shows the position of the PCR primers.

**(B)** Top: Schematic (not to scale) of the plasmids used to delete different regions of the *pfap2-g* locus using an alternative three plasmids strategy. Bottom: Diagnostic PCRs to assess correct edition in the transgenic lines generated with this strategy and their subclones (subclones were obtained to achieve pure populations of edited parasites).

**Table S1. Oligonucleotide sequences.** List of the oligonucleotides used in the study, classified according to their use.

| Oligonucelotides used for 5'RACE | |
|---|---|
| **Name** | **Sequence (5'-3')** |
| 5'RACE_RNA_adapter | GCUGAUGGCGAUGAAUGAACACUGCGUUUGCUGGC |
| 5'RACE_Outer | GATGGCGATGAATGAACACTG |
| Pfap2-g_+345_R | GATACATTCTCGTTACTCTGCA |
| 5'RACE_Inner | GAATGAACACTGCGTTTGCTG |
| Pfap2-g_+272_R | CATGCTCTCTTCCCATTCGAA |

| Primers for cloning steps | |
|---|---|
| **Name** | **Sequence (5'-3')** |
| Pfap2g_-3929_HR1_KO_F_NotI | tggtgtgcggccgcCGTACATATATACAAATAGAGAT |
| Pfap2g_-3521_HR1_KO_R_PstI | tggtgtctgcagACAGGTATTGTACGCCTTTTA |
| Pfap2g_+991_HR2_KO_F_SpeI | tgttggactagtCCTTGAAAAGAATATAGAAGAAC |
| Pfap2g_+1973_HR2_KO_R_AflII | tggtgtcttaagCAATTTATCAGCATCGTCATCA |
| GFP_+3_F_PstI | tggtgtctgcagAGTAAAGGAGAAGAACTTTTCA |
| GFP_+714_R_SpeI | tggtgtactagtTTATTTGTATAGTTCATCCATGC |
| Pfap2g_-3503_Guide_F | taagtatataatattTTCTTTTTAAGGTTGCGTACgttttagagctagaa |
| Pfap2g_-3522_Guide_R | ttctagctctaaaacGTACGCAACCTTAAAAAGAAaatattatatactta |
| Pfap2-g_+940_Guide_F | taagtatataatattTCATAAGAAATATGATTCATgttttagagctagaa |
| Pfap2-g_+921_Guide_R | ttctagctctaaaacATGAATCATATTTCTTATGAaatattatatactta |
| 1144400_+1144_HR1_F_SacII | tggccgcggGGAAAGACAATCAACAAACCTT |
| 1144400_+1562_HR1_R_SpeINcoI | tggtgtactagtctgccatggTCACTTGAAATCGTTTTGTTATTA |
| 1144400_+1646_HR2_F_SpeI | tggttgactagtATTGTATAAATGAAGAGACATACA |
| 1144400_+2199_HR2_R_AflII | tggtgtcttaagTGATTATTATGTGTAACTTCAGTA |
| ap2g_-181_F1_F_NcoI | tggtgtccatggGTAGGTACATTCAAATATCTCC |
| ap2g_+992_F1_R_SpeI | tggtgtactagtTTCTTCTATATTCTTTTCAAGGAT |
| ap2g_-1200_F2_F_NcoI | tggtgtccatggGATACTTTATGATTAGTTGATATG |
| ap2g_+118_F2_R_SpeI | tggagaactagtTTCCTGGGATGTAATCAAAAGT |
| ap2g_-2147_F3_F_NcoI | tggtgtccatggATAATGCTCTGAGAATATATACTT |
| ap2g_-891_F3_R_SpeI | tgttggactagtAATTTATATATCTCGACACCTTTA |
| ap2g_-3000_F4_F_NcoI | tggtgtccatggTATTGACGTGCATACTATATGTA |
| ap2g_-1834_F4_R_SpeI | tggagaactagtAATGTATTTCATGATATTAGTGTAA |
| ama1_-966_F_NcoI | tggtgtccatggGTTATGTGTAGAATATTTACAAAG |
| ama1_+249_R_SpeI | tgttggactagtACCTTCGTGGTCTATTGGATA |
| Pfap2-g_+425_nucleo_R_SpeI | tggtgtactagtTCTATGATTTAACAACATATCCAA |
| Pfap2-g_-6_2nucleo_F_NcoI | tggtgtccatggAAGAAGATGACCGCTAAGATAT |
| Pfap2-g_+210_1nucleo_F_NcoI | tggtgtccatggATTGAAGAAAGTTGTTTTGACATT |
| Pfap2-g_+341_RestF1_F_NcoI | tggtgtccatggTTGCAGAGTAACGAGAATGTAT |
| Pfap2-g_+748_F0_F_NcoI | tggtgtccatggGAAATGTAAAAGAGTATTCTCATA |
| Pfap2-g_+1951_F0_R_SpeI | tggtgtactagtCAATTTATCAGCATCGTCATCA |
| 1144400_+1566_Guide_F | taagtatataatattATTATGTGTTCCAAGAACGAgttttagagctagaa |
| 1144400_+1585_Guide_R | ttctagctctaaaacTCGTTCTTGGAACACATAATaatattatatactta |
| PFL1950W_-1096_F_NcoI | tggtgtccatggGACATTAGTTATATGAATATATTGG |
| PFL1950W_+112_R_SpeI | acaccaactagtGCATCTCTTTGTACTTCTGCG |
| AMA1_-1692_F_NcoI | tggtgtccatggTATACGAGTAAAACACATCACAG |
| AMA1_+250_R_SpeI | acaccaactagtGTTCTTGTGGTGCGGGTTCG |
| mspdbl2_+705_NcoI_F | tggtgtccatggTCATATGCATCATCTGAAGC |
| mspdbl2_+1883_SpeI_R | acaccaactagtTCTATTTCCCCATCTGTATTA |
| mspdbl2_-173_NcoI_F | tggtgtccatggTATTTGTTCATTGAACTGTTAATG |
| mspdbl2_+1004_SpeI_R | acaccaactagtGTACATTGGTTATCTTTCTGC |
| mspdbl2_-1013_NcoI_F | tggtgtccatggGATAAGTCGCTGCGATTCC |
| mspdbl2_+120_SpeI_R | acaccaactagtTATGTTATTTCTTAAATTAGGGT |
| varB_-1412_NcoI_F | tggtgtccatggCAAGGTAATTTCATACATATGTG |
| 0733000_-1_speI_R | acaccaactagtTTTCGTTATTGGTGCACTACAT |
| 1041300_-1_speI_R | acaccaactagtTGTACCGACAACATGATGTTAT |
| 1041300_-1096_F_NcoI | tggtgtccatggTGATTGAAGAATACGTATGCCT |
| 1041300_+148_R_SpeI | acaccaactagtACAAACGTCCATGCAATTGAC |
| Pfap2-g_-448_HR1_KOF1_F | cggtacccggggatccATATGTCCTATAGGTGTCAAAC |
| Pfap2-g_-159_HR1_KOF1_R | tcttctcctttactctgcagGGAGATATTTGAATGTACCTAC |
| Pfap2-g_+993_HR2_KOF1_F | atacaaataaactagtCCTTGAAAAGAATATAGAAGAAC |
| Pfap2-g_+1974_HR2_KOF1_R | gactctagaggatccCAATTTATCAGCATCGTCATCA |
| Pfap2-g_-159_Guide_F | ttctagctctaaaacTTATATTGGCACTAATTTAGaatattatatactta |
| Pfap2-g_-139_Guide_R | taagtatataatattCTAAATTAGTGCCAATATAAgttttagagctagaa |

| Primers for cloning steps | |
|---|---|
| **Name** | **Sequence (5'-3')** |
| Pfap2-g_-1857_HR1_KOF2-3_F | cggtacccggggatccTTACACTAATATCATGAAATACATT |
| Pfap2-g_-1427_HR1_KOF2-3_R | tcttctcctttactctgcagACAAAATCTATAATCTTATATATGAA |
| Pfap2-g_-129_HR2_KOF2-3_F | atacaaataaactagtAATTTGAAGTACCAACATATACAT |
| Pfap2-g_+234_HR2_KOF2-3_R | gactctagaggatccAATGTCAAAACAACTTTCTTCAAT |
| Pfap2-g_-1412_Guide_F | taagtatataatattTTTAATAATACGTATGCTTGgttttagagctagaa |
| Pfap2-g_-1393_Guide_R | ttctagctctaaaacCAAGCATACGTATTATTAAAaatattatatactta |
| ap2g_-479_InF_SacII_F | actaaatatatatccaatggccgcggCTTTAATGTTGTATGTATGTTTA |
| ap2g_-1_InF_SpeI_HindIII_R | actagtacacaccaaagcttCTTCTTAAATATTCCTTATTAAAA |
| ap2g_+993_InF_HindIII_SpeI_F | aagctttggtgtgtactagtCCTTGAAAAGAATATAGAAGAAC |
| ap2g_+1974_InF_EcoRI_R | ttccccgaaaagtgccacctgaattcCAATTTATCAGCATCGTCATCA |
| ap2g_-1402_InF_SacII_F | actaaatatatatccaatggccgcggCGTATGCTTGTGGATATTATAA |
| ap2g_-900_InF_HindIII_AflII_R | aagcttacacaccacttaagTCTCGACACCTTTATTATATATA |
| ap2g_+1493_InF_AflII_HindIII_F | cttaagtggtgtgtaagcttATAAGAAGAACAATAAAAATAAGA |
| ap2g_+1981_InF_NcoI_R | gtgttattattttaccgttccatggTATTATCCAATTTATCAGCATC |
| ap2g_-3929_InF_SacII_F | actaaatatatatccaatggccgcggCGTACATATATACAAATAGAGAT |
| ap2g_-3522_InF_HindIII_AflII_R | aagcttacacaccacttaagACAGGTATTGTACGCCTTTTA |
| ap2g_+1_InF_AflII_HindIII_F | cttaagtggtgtgtaagcttATGACCGCTAAGATATTAAATC |
| ap2g_+521_InF_NcoI_R | gtgttattattttaccgttccatggATATACGCTCATTTCCTTTATC |
| ap2g_+493_InF_SacII_F | actaaatatatatccaatggccgcggAAAGAGGATAAAGGAAATGAG |
| ap2g_+984_InF_HindIII_AflII_R | aagcttacacaccacttaagCATATTACCTATATAAGTTTTAG |
| ap2g_+7299_InF_AflII_HindIII_F | cttaagtggtgtgtaagcttATATCATCCTTTTTTTTTATAAAAT |
| ap2g_+7744_InF_NcoI_R | gtgttattattttaccgttccatggACCAAGTTCATTTTAATGTTATT |
| GFP_+3_AflII_F | tggtgtcttaagAGTAAAGGAGAAGAACTTTTC |
| GFP_+714_HindIII_R | acaccaaagcttTTATTTGTATAGTTCATCCATG |
| GFP_+3_HindIII_F | tggtgtaagcttAGTAAAGGAGAAGAACTTTTC |
| GFP_+714_SpeI_R | acaccaactagtTTATTTGTATAGTTCATCCATG |
| ap2g_+45_guide1_F | taagtatataatattAGGGTATACAGGGATATCGAgttttagagctagaa |
| ap2g_+45_guide1_R | ttctagctctaaaacTCGATATCCCTGTATACCCTaatattatatactta |
| ap2g_-827_guide1_F | taagtatataatattTATGATATCAAATTAAAATAgttttagagctagaa |
| ap2g_-827_guide1_R | ttctagctctaaaacTATTTTAATTTGATATCATAaatattatatactta |
| ap2g_+1453_guide2_F | taagtatataatattAAAGATGAACACAAGAAGGAgttttagagctagaa |
| ap2g_+1453_guide2_R | ttctagctctaaaacTCCTTCTTGTGTTCATCTTTaatattatatactta |
| ap2g_-199_guide2_F | taagtatataatattGTAACTATCTATATATAAGTgttttagagctagaa |
| ap2g_-199_guide2_R | ttctagctctaaaacACTTATATATAGATAGTTACaatattatatactta |
| ap2g_+1104_guide1_F | taagtatataatattTTATAGTAATCAAACATGCAgttttagagctagaa |
| ap2g_+1104_guide1_R | ttctagctctaaaacTGCATGTTTGATTACTATAAaatattatatactta |
| ap2g_-1427_InF_HindIII_AflII_R | aagcttacacaccacttaagACAAAATCTATAATCTTATATATGAA |
| ap2g_-130_InF_AflII_HindIII_F | cttaagtggtgtgtaagcttAATTTGAAGTACCAACATATACAT |
| ap2g_+234_InF_NcoI_R | gtgttattattttaccgttccatggAATGTCAAAACAACTTTCTTCAAT |
| ap2g_-1633_gRNA_R | taagtatataatattAAAAGTATTTATATCTCTCAgttttagagctagaa |
| ap2g_-1633_gRNA_F | ttctagctctaaaacTGAGAGATATAAATACTTTTaatattatatactta |
| ap2g_-1402_InF_AflII_HindIII_F | cttaagtggtgtgtaagcttCGTATGCTTGTGGATATTATAA |
| ap2g_-900_InF_NcoI_R | gtgttattattttaccgttccatggTCTCGACACCTTTATTATATATA |

| Primers for diagnostic PCR | |
|---|---|
| **Name** | **Sequence (5'-3')** |
| 1144400_+865_F | GATATTATGAGTACTGATAGTGA |
| 1144400_+2319_R | TGACTCGTTCAGGTTATTCTTA |
| 1144400_+1799_R | CAGATCTATATTTCATTTGTTCTT |
| Pfap2-g_-2147_UpTSS1_F | TAATGCTCTGAGAATATATACTTA |
| Pfap2-g_UpTSSR_-710 | TGTATGTAAAAATAAAGCACCAC |
| Pfap2-g_CDS_+345_R | GATACATTCTCGTTACTCTGC |
| Pfap2-g_5'UTR_-460_F | GCTTCTTTAATGTTGTATGTATG |
| Pfap2-g _+2474_R | TTACTCCATTAGGTGCATTCAT |
| 1144300_+4_F | CACATGGAGCAAGCAGGTAT |
| 1144300_+632_R | TTTTGTAAACGTCTGGTTCTCT |
| 1144300_UTR_F | TAACAAATATGAATATTGAAGGATA |
| 1144400_-904_F | TAAGTATGGTTCATTTAGGGGT |
| 1144400_+22_R | TGTCGTCTTTAATCAGTTCCAT |
| Pfap2-g_UpTSSF_ -1820 | ATGGCTTTTATTTATTCTTAATTGT |
| Ap2g_3' EXT KO1KO2_R_+2277 | CAAACGGTTACTATTATTACTATT |
| Ap2g_5' EXT KO3_F_-4300 | GAAAATTAAACATTGGAGAATTGA |
| Ap2g_3' EXT KO3_R_+755 | CATTTCATCAAAGACGCCGTA |
| Ap2g_5' EXT KO4_F_+200 | GTATGAAATAATTGAAGAAAGTTG |
| Ap2g_3' EXT KO4_R_+8133 | GTACAACAAAAACTGAACTCTTT |

| Primers for gDNA qPCR and ChIP-qPCR | |
|---|---|
| **Name** | **Sequence (5'-3')** |
| Pfap2-g_-2147_UpTSS1_F | TAATGCTCTGAGAATATATACTTA |
| Pfap2-g_-1979_UpTSS1_R | CAATTAGGATAAATAAATATTCCAT |
| Pfap2-g_ -1820_UpTSS2_F | ATGGCTTTTATTTATTCTTAATTGT |
| Pfap2-g_ -1644_UpTSS2_R | TATCTCTCACGGTTCATATCTT |
| Pfap2-g_-1409_DwnTSS_F | TAATACGTATGCTTGTGGATATT |
| Pfap2-g_-1226_DwnTSS_R | ATATGGAAACTAATAATAAATTGTTA |
| PF3D7_1222600_F1 (CDS) | AACAACGTTCATTCATTCAATAAATAAGG |
| PF3D7_1222600_R1 (CDS) | ATGTTAATGTTCCCAAACAACCG |
| Seryl_qPCR_F (serrs) | AAGTAGCAGGTCATCGTGGTT |
| Seryl_qPCR_R (serrs) | TTCGGCACATTCTTCCATAA |
| hdhfr_F | AGTAGAAGGTAAACAGAATCT |
| hdhfr_R | GGCATCATCTAGACTTCTGG |
| 5'HR_1144400_+1424_F | TAATGCAAAACGATAGTAATCTTA |
| 5'HR_1144400_+1526_R | TCTTCATTAGGATCATATCCATT |
| 3'HR_1144400_+1676_F | TGAAACTTGAAAACTGTGATGAA |
| 3'HR_1144400_+1799_R | CAGATCTATATTTCATTTGTTCTT |
| PFL1950wF | CTATGTTGTATTATTCGATATTTTC |
| PFL1950wR | AGAATAGGAAAATACAAATTATAGC |
| Pfap2-g_CDS_+259F | GAAGAGAGCATGCAATGAAGT |
| Pfap2-g_CDS_+382R | TTGTCCATGCAACTATTCGATA |
| Pfap2-g_-1409_DwnTSS_F | TAATACGTATGCTTGTGGATATT |
| Pfap2-g_-1226_DwnTSS_R | ATATGGAAACTAATAATAAATTGTTA |
| Ap2-g_-2967_F | ATTATTACCTTCGGTACCTTAAT |
| Ap2-g_-2861_R | AATGCACTTTTTGAGTACAGTTA |
| PFL1085w_-2553_A2.5F | CAATATAATACCATACTTCAAAAC |
| PFL1085w_-2460_A2.5R | AGTATGGTTTTGTACTCCTTTTA |
| AMA1_+23_F | ATTATTGAGCGCCTTTGAGTTT |
| AMA1_+145_R | GTGTAATGGATATTCGTATTCTT |
| Pfap2-g_+653_F | TTTCCCATACTATATGCTGAAAT |
| Pfap2-g_+774_R | CATATGAGAATACTCTTTTACATT |
| Pfap2-g_+754_F | GTAAAAGAGTATTCTCATATGTAA |
| Pfap2-g_+876_R | AACATAAAAGTACATGACATCATT |
| clag3.2_5_F | TAGGCGAAAATAAAAACGAAAATG |
| clag3.1_clag3.2_5_R | CATGGATTTTAATTGTTCAATATTG |
| 5'ext_1144400_+734_F | ATGAAGTAGCTACTACAATACAT |
| 5'ext_1144400_+870_R | AATATCCATAAGAACATCCTTGT |
| 3'ext_1144400_+2177_F | CTGAAGTTACACATAATAATCAAA |
| 3'ext_1144400_+2319_R | TGACTCGTTCAGGTTATTCTTA |
| mspdbl2_-697_F | TCATGTATATGTAAAAGTGATTCT |
| mspdbl2_-482_R | CCCTCAAATAAAGTGCATTATG |
| mspdbl2_+766_F | CTCATCAAGCTATAAGATATAGT |
| mspdbl2_+966_R | CATCATTGCTTCCCAAACATG |
| Pfap2-g_-2147_UpTSS1_F | TAATGCTCTGAGAATATATACTTA |
| Pfap2-g_-1979_UpTSS1_R | CAATTAGGATAAATAAATATTCCAT |
| PF3D7_1222600_F1 | AACAACGTTCATTCATTCAATAAATAAGG |
| PF3D7_1222600_R1 | ATGTTAATGTTCCCAAACAACCG |
| 1041300_-129_F | TCTACCATATTACAATACTCCC |
| 1041300_-9_R | ACATGATGTTATACAATTTGGTG |
| 0733000_-1013_F | TGGTATCAAGAATACGTATGCT |
| 0733000_-842_R | GAACAAGAACGTATCGATATAG |
| Ap2g_3' EXT KO1KO2_F _+2088 | CAACCTAACGTTCTTGAAAAAG |
| Ap2g_3' EXT KO2KO2_R_+2277 | CAAACGGTTACTATTATTACTATT |
| Ap2g_5' EXT KO3_F_-4300 | GAAAATTAAACATTGGAGAATTGA |
| Ap2g_5' EXT KO3_R_-4138 | TTACATATACATATACATACATTTA |
| Ap2g_HR1 KO3_F_-3902 | GCTAGACGTACAATAATATACAA |
| Ap2g_HR1 KO3_R_-3552 | TTTTTTATTATTTGCTATTTTTAAAG |
| ap2g_-805_R | GAAACCTTTTGAATTTATAATATTAT |
| ap2g_-1547_F | GATAAAGTAGAATATTGTTCTATAT |
| ap2g_-1392_R | TGCATATGTATAAATATTGTCAGT |
| ap2g_-3760_F | GCACATATATTTTAATTAGCTAGA |
| ap2g_-3708_F | CTTTATATATTGTATTTATGCATAT |
| ap2g_-3757_R | GTGCACAAGATTAATAATAATTAT |
| ap2g_-3523_R | ACAGGTATTGTACGCCTTTTAA |

| Primers for gDNA qPCR and ChIP-qPCR | |
|---|---|
| **Name** | **Sequence (5'-3')** |
| ap2g_-5004_F | CTGAATATTTTTCTTTTGGTCGT |
| ap2g_-4861_R | GACATAATATATATTGAGCATGC |
| ap2g_-4359_F | AATAGATAAAGTAGAATATTGTTCT |
| ap2g_-4200_R | GCATATGTATAAATATTGTCAGTT |
| ap2g_-3930_F | CGTACATATATACAAATAGAGATT |
| ap2g_-3737_R | TCTAGCTAATTAAAATATATGTGC |
| Pfap2-g_UpTSSF_ -1820 | ATGGCTTTTATTTATTCTTAATTGT |
| ap2g_-1321_F | GAATGTTGTAGTTCTTTCTTTCT |
| ap2g_-1177_R | CATATCAACTAATCATAAAGTATC |
| Pfap2-g_qPCR_-563_R | TGCAAAAACATGCACATAATATG |
| Pfap2-g_qPCR_-484_F | GCTTCTTTAATGTTGTATGTATG |
| Pfap2-g_qPCR_-327_R | GTTAGTAATACAAAAATGTAAATAC |
| ap2g_-352_F | GTATTTACATTTTTGTATTACTAAC |
| ap2g_-160_R | GGAGATATTTGAATGTACCTAC |
| Ap2g_5' EXT KO4_F_+200 | GTATGAAATAATTGAAGAAAGTTG |
| Pfap2-g_CDS_+345_R | GATACATTCTCGTTACTCTGC |
| Ap2g_3' EXT KO3_F_+618 | CATCTATTTATTGTCAAGAAAGAA |
| Ap2g_3' EXT KO3_R_+755 | CATTTCATCAAAGACGCCGTA |
| PfAP2-G +1106 F | TAGTAATCAAACATGCATGGATT |
| PfAP2-G +1237 R | CAACTACAGGTACAGAACATAT |
| Ap2g_HR2 KO2_F_+1721 | TTCTCATTCTTCAAATAGCCTTG |
| Ap2g_HR2 KO2_R_+1866 | TTTAACATTATCTTTGTGAGAATAA |
| ap2g_+2017_IPC_F | GATTATTTTACTATTTGTGATCCT |
| ap2g_+2183_IPC_R | TGATTACTTATAGAATTATGCTTG |
| pfap2-g_+3979_R | ATGTTAATGTTCCCAAACAACCG |
| pfap2-g_+3874_F | AACAACGTTCATTCAATAAATAAGG |
| PFL1085w_F2 | GCTACCTTACTAATGTATTAAGTC |
| PFL1085w_R2 | CATATTATTAAAGAATTGGTTCACG |
| PFL1085w_+7357_F3 | CCTTTTGAATATTCCCTATAACT |
| PFL1085w_+7467_R3 | GTTATTTAAAAATGTCACAAGATG |
| Ap2g_3' EXT KO4_F_+7981 | TTTTCCACATTATAAGAACTGTAT |
| Ap2g_3' EXT KO4_R_+8133 | GTACAACAAAAACTGAACTCTTT |
| ap2g_+26_F | TCGATATCCCTGTATACCCTT |
| ap2g_+224_R | CAACTTTCTTCAATTATTTCATAC |
| GFP_+1_F | GTAAAGGAGAAGAACTTTTCAC |
| GFP_+158_R | GGTAGTTTTCCAGTAGTGCAA |
| GFP_+580_F | TTACCAGACAACCATTACCTG |
| GFP_+714_R | TTATTTGTATAGTTCATCCATGC |
| GFP_+348 _F | GTGATACCCTTGTTAATAGAAT |
| GFP_+540_R | ATGGTCTGCTAGTTGAACGC |
